# Molecular Photoswitches Regulating the Activity of the Human Serotonin Transporter

**DOI:** 10.1101/2023.09.20.558680

**Authors:** Nadja K. Singer, Leticia González, Antonio Monari

## Abstract

Serotonin is an essential mediator regulating diverse neural processes, and its deregulation is related to debilitating neurological diseases. In particular, the human serotonin transporter (hSERT) is fundamental in completing the synaptic neural cycle by allowing the reuptake of serotonin. Its inhibition is particularly attractive, especially as a pharmacological target against depressive syndrome. Here, we analyze, by using long-range molecular dynamic simulations, the behavior of a molecular photoswitch whose *cis*- and *trans*-isomers inhibit the hSERT differently. In particular, we evidence the structural and molecular basis behind the higher inhibiting capacity of the *cis*-isomer, which blocks more efficiently the hSERT conformational cycle leading to serotonin uptake.

**TOC Graphic:** 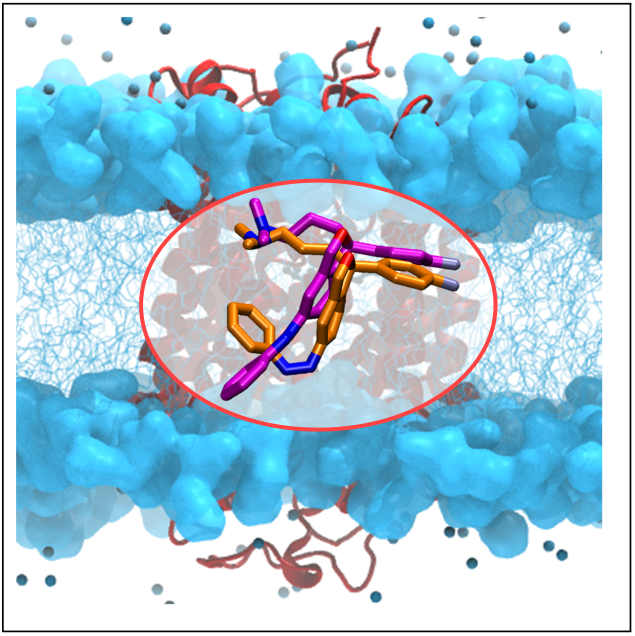

---

The human serotonin transporter (hSERT) is a membrane protein, which plays an essential role in the modulation of neurotransmission. Indeed, the hSERT is a transmebrane transporter allowing the sodium- and chloride-dependent re-uptake (Fig. 1A) of the neurotransmitter serotonin (5-hydroxytryptamin, 5HT, Fig. 1B) from the synaptic cleft into the presynaptic neurons, thus terminating the serotonogenic signaling. Serotonin re-uptake proceeds through a conformational cycle in which the hSERT goes from the outward-open conformation, exposing the 5HT binding pocket to the external medium, back to the inward-open conformation, which allows release of 5HT inside the neuronal cellular compartments (Fig. 1A). The precise regulation of the underlying conformational transitions is thus fundamental in dictating the overall hSERT efficiency and its regulatory role. Dysfunctional or improper regulation of neurotransmitter transporters, including the hSERT, have been associated with highly debilitating neurological and neuropsychiatric disorders, such as depression, attention-deficit hyperactivity, epilepsy, and narcolepsy.^1–3^ Therefore, the hSERT has emerged as a key target for the development of drugs aimed at managing major neuropsychiatric conditions. As a most relevant example, the antidepressant citalopram (Fig. 1B) is a clinically approved drug inhibiting the hSERT. It belongs to a larger family known as selective serotonin reuptake inhibitors (SSRIs).^4^ Interestingly, citalopram was initially commercialized by the pharmaceutical company Lundbeck as a racemic mixture, however, today only the more potent escitalopram (i.e. (*S*)-citalopram, Fig. 1B) is used. The mechanism of action underlying the biological activity of escitalopram has been characterized, also from a biophysical and structural point of view, by protein X-Ray crystallography.^5^ It has been recognized that the drug binds to the orthosteric serotonin binding site of the hSERT and subsequently locks the transporter in an outward-open conformation (Fig. 1A), hence directly inhibiting serotonin re-uptake. Additionally, escitalopram was also found to bind to the hSERT allosteric binding site, which represents a gateway to the orthosteric binding site, thus offering a secondary inhibitory mechanism and ultimately resulting in one transporter being, potentially, blocked by two drug molecules.^5^ As a matter of fact, escitalopram’s binding efficiency and flexibility clearly makes it an extremely interesting scaffold for further drug development. As an example, hSERT-inhibiting derivatives of escitalopram have been recently published by Cheng et al. ^6^ and Dreier.^7^ Interestingly, in these contributions the escitalopram scaffold has been combined with a photoswitching unit, which allows to modulate the inhibiting efficiency of the drug by the use of a suitable light perturbation. Because of the role of the hSERT in modulating different neurological functions, the possibility to dispose of photo-activable inhibitors paves the way to most suitable optogenetic-like applications. Indeed, by combining escitalopram with an azo-benzene unit, Cheng et al.^6^ obtained a molecule that can be converted from the thermodynamically more stable *trans*-to the less stable *cis*-azo-escitalopram through irradiation with UV light at 365 nm (see Fig. 1B). The backward *cis*-to *trans*-azo-escitalopram conversion is achieved by irradiation at 460 nm. Interestingly, the two isomers present a very different inhibition efficiency, with *cis*-azo-escitalopram, being *∼*43-fold more active against the hSERT,^6^ thus offering an efficient way to control serotonin uptake through an external light stimulus.

**Figure 1.**
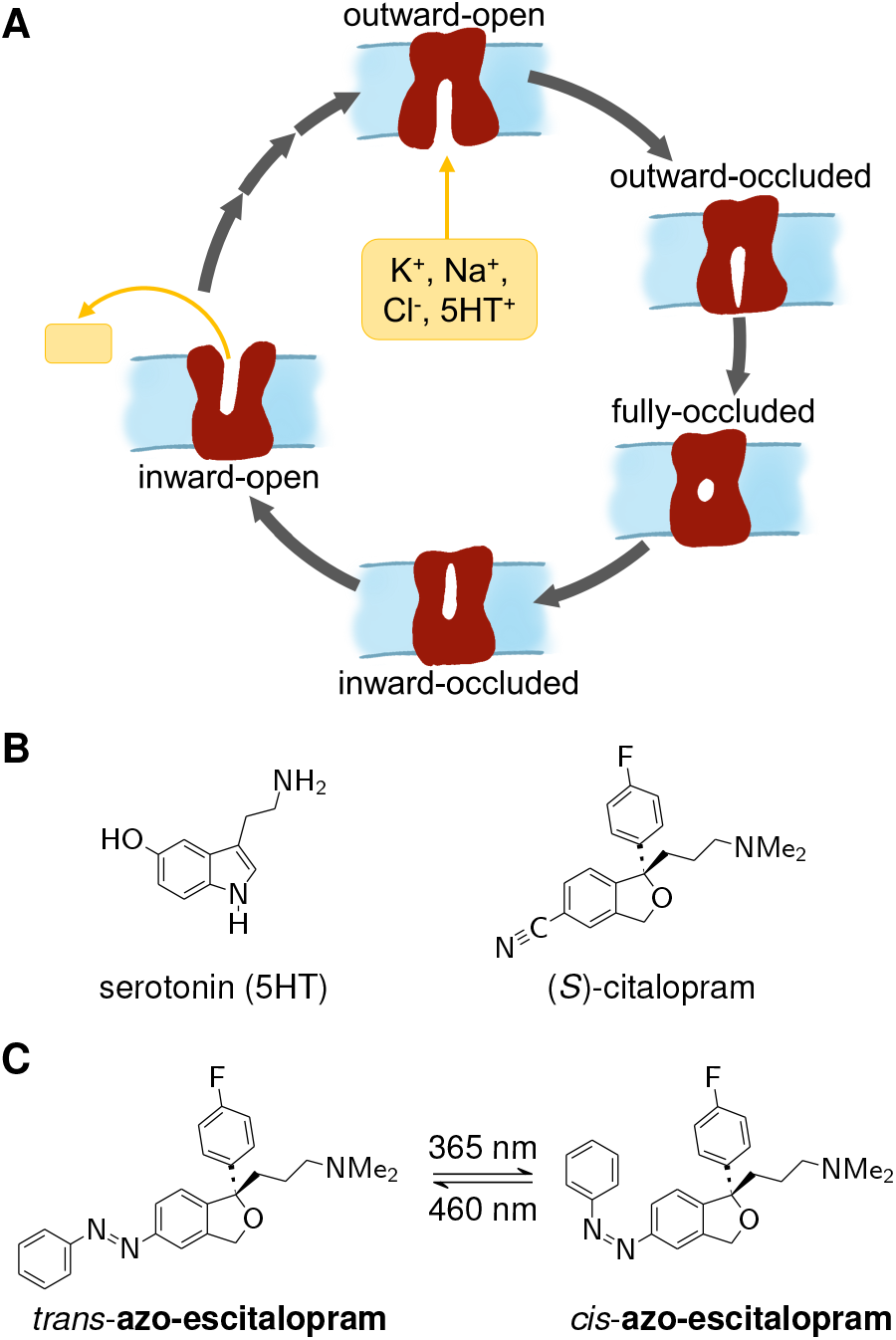
A) Schematic representation of the key steps (outward-open, outward-occluded, fully-occluded, inward-occluded, inward-open) in the physiological transport cycle of the hSERT. B) Scheme of the *trans*- and *cis*-azo-escitalopram photoswitch.

The use of photoswitches is growing in different scientific and technological fields, ^8^ including molecular machines, smart electronics,^9^ and more recently photopharmacology.^10^ The latter field also involves the use of biomimetic photoswitches to selectively induce cellular membrane destabilization and hence cytotoxicity.^11–13^ For instance, the study of retinal-based photoswitches in different classes of rhodopsin has led to significant advances in understanding fundamental photophysical processes, including the rationalization of signal transduction in vision,^14^ and the development of optogenetic applications.^15^ In contrast, the study of photoswitches that can modulate the synaptic signal-transduction and in the hSERT is less mature. For example, despite a different behavior having been evidenced experimentally,^6^ the molecular basis underlying the differential inhibiting power of the two photoisomers of azo-escitalopram is far from being properly characterized and understood. In this Letter, we resort to long-scale classical molecular dynamics (MD) simulations and binding free energy estimations for the hSERT embedded in a model lipid bilayer and interacting with *trans*- and *cis*-azo-escitalopram. This strategy will provide an atomistic level description of the perturbation of the protein structure induced by the ligand, as well as the intrinsic binding affinity, thus allowing to formulate hypotheses on the factors ultimately modulating the inhibition efficiency.

After a preliminary conformational search done using CREST 2.10.2,^16,17^ we employed Density Functional Theory (DFT) to optimize the ground state geometries of both azo-escitalopram isomers (Fig. 1B) at the CAM-B3LYP-D3/def2-TZVP^18–20^ level of theory. The electronic structure calculations have been performed using Gaussian 16.^21^ Gibbs free energies at 298.15 K and the optimized XYZ-coordinates are reported in Section S1 of the Supporting Information.

Subsequently, we built an initial model of the hSERT from the outward-open X-ray structure (PDB entry 5I71 chain A, referred to hereafter as 5I71) resolved by Coleman et al.,^5^ where the mutations I291A, T439S, and Y110A were reverted back to the native amino acids with the SwissModel server.^22^ The reconstructed hSERT was embedded into an explicit 1-palmitoyl-2-oleoyl-sn-glycero-3-phosphatidylcholine (POPC) lipid bilayer involving 175 lipids on the lower leaflet and 173 in the upper layer, using the CHARMM-GUI input generator,^23–26^ after reorientation with the OPM server,^27^ and explicitly including the C200-C209 disulfide bond. Furthermore, residues E508 and E143 were set to their *ϵ*-nitrogen protonated forms.^28^ Structural water molecules and ions (2xNa^+^, 1xCl^*−*^) in the binding site are essential for substrate transport and ligand interaction and hence should be properly modeled. Since 5I71 did not include the chloride ion the position of the latter was inferred from the hSERT PDB entry 5I6X (chain A, referred to hereafter as 5I6X).^5^ Finally, the system was solvated with a water buffer of 40 Å including salt (KCl, NaCl) at a physiological concentration of 0.15 M. The ligands were placed in the orthosteric binding site of the hSERT by aligning them to the escitalopram scaffold in 5I71, as well as through docking with HDock.^29^ The final system contained *∼*150,000 atoms including 348 lipids, 92 Na^+^, 89 K^+^, and 184 Cl^*−*^ ions as well as 33,506 water molecules with a box volume of *∼*12 nm^3^. In addition, systems of *trans*- and *cis*-azo-escitalopram in water were prepared, each containing *∼*75,000 atoms with a box volume of *∼*9.5 nm^3^. Equilibrium MD simulations have been run using the NAMD code,^30,31^ the lipid and protein being represented by Amber’s Lipid17^32^ and ff14SB^33^ force fields, respectively. Water was modeled with TIP3P^34^ and the ion parameters of Li/Merz^35^ were used. The photoactive drug force field was obtained following the generalized amber force field (GAFF) approach as implemented in AmberTools.^36^

Isobaric and isothermal conditions (NPT) have been applied, and Newton’s equations of motion have been integrated with a time step of 4 fs using Hydrogen Mass Repartition^37^ in combination with Rattle^38^ and Shake^39^ at 300 K and 1 atm. Before the production runs, minimization and equilibration have been performed progressively relaxing the harmonic constrains on the protein backbone, lipids, and ligands. All the analysis (see also Section S2) have been performed using NAMD^30,31^ and VMD^40^ utilities.

The DFT geometry optimization of the two isomers is coherent with the well-known behavior of azobenzene derivatives, and while the azobenzene unit in the *trans*-isomer is largely planar, the sterical hindrance of the *cis*-counterpart, flips one phenyl ring of the azobenzene unit out of plane (similar to Fig. 2C). The Gibbs free energy, estimated from the harmonic vibrational frequencies and the rigid rotor approximation, indicates, similarly to the experimental data, that the *trans*-isomer is thermodynamically favored by about 11 kcal mol^*−*1^.

**Figure 2.**
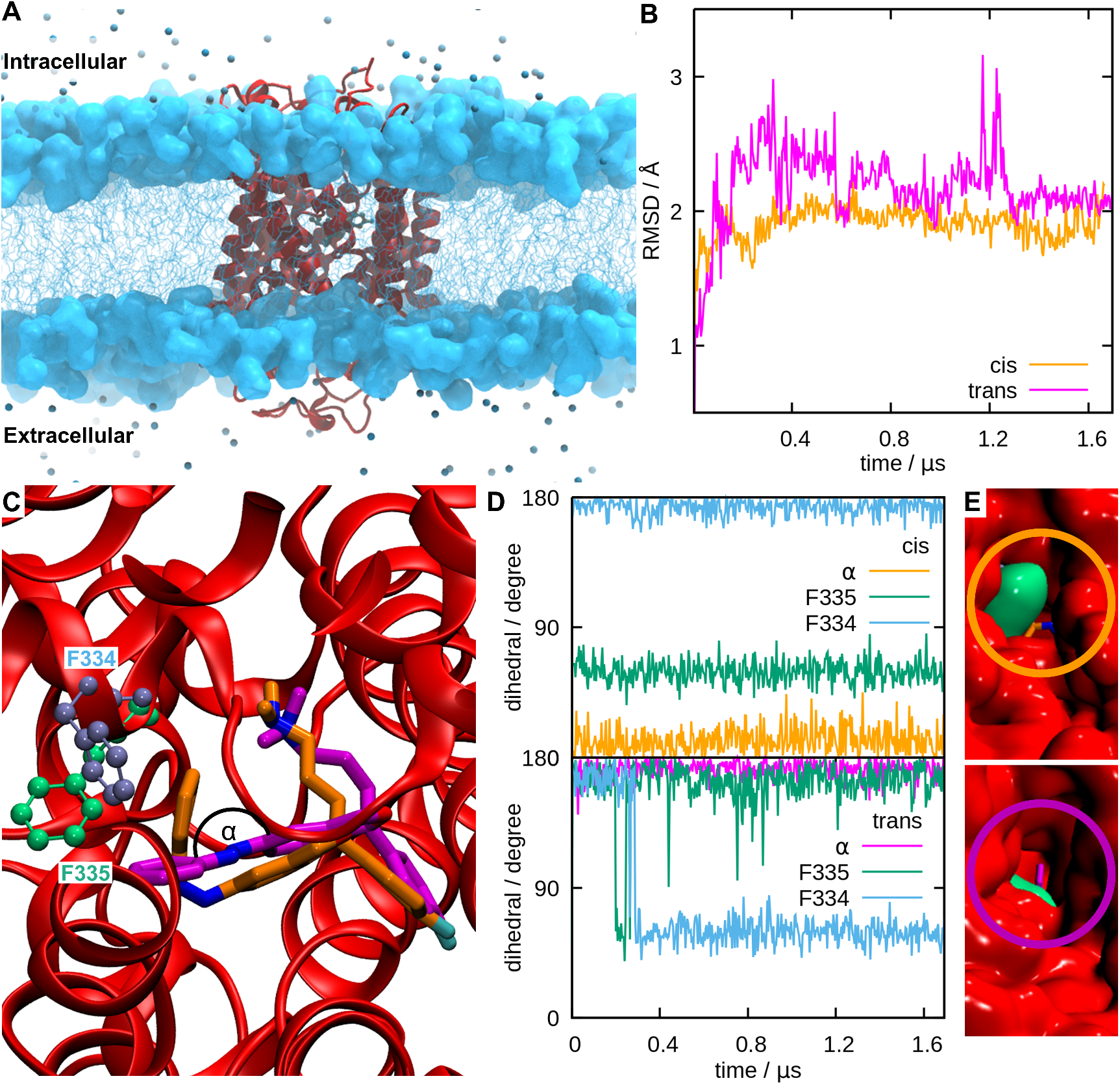
A) Representative snapshot of the hSERT from the MD simulations. B) Time series of the hSERT C*α*’s RMSD. *trans*-(pink) and *cis*-azo-escitalopram (orange) are shown in reference to their starting structure (frame 0). C) Zoom into the central escitalopram binding site with F334 and F335, as well as the azo-(CNNC) dihedral angle *α* highlighted. D) Time series of the CNNC dihedral angle *α* of the azo bond of azo-escitalopram, F334, and F335 along the *cis*-(top) and *trans*-azo-escitalopram simulation (bottom). E) View from the extracellular side of *cis*-(top) and *trans*-azo-escitalopram (bottom) buried by residue F335.

Next, we resort to studying the differential interaction modes of the two isomers with the hSERT. A representative snapshot of the full system embedded in the lipid bilayer is reported in Figure 2A. Equilibrium MD simulations have confirmed that both azo-escitalopram isomers bind persistently in the orthosteric site of the hSERT. In fact, not only no spontaneous release of the ligand is observed, but the global structure of the transporter remains stable, showing only moderate oscillations, which can be appreciated from the time series of the root mean square deviation (RMSD) reported in Figure 2B. Furthermore, the RMSD time-series do not exceed 3 Å and reach a plateau at the end of our simulation. Interestingly, even if the RMSD remains globally modest, a slightly higher rigidity can be observed when the *cis*-isomer is present, suggesting possible differences in the specific binding modes, and a stronger locking propensity of the latter. Representative snapshots that superpose the binding poses of the two isomers can be seen in Figure 2C. As expected, while the global features of the escitalopram moiety interacting with hSERT are conserved among the two isomers, some differences can be evidenced. In fact, the planar and rigid azobenzene moiety of the *trans*-isomer induces additional constraints compared to the more globular *cis*-conformation. In turn, this is translated in a deeper buried escitalopram unit for the *trans*-configuration (Fig. 2C). Furthermore, a differential interaction pathway, also due to potential sterical clashes, is evidenced for the azobenzene moiety and F334 and F335 amino acids, as it is highlighted in Figure 2C. This different interaction can also be observed by monitoring the time series of the dihedral angles defining the orientation of the two amino acid lateral chains with respect to the backbone (Fig. 2D). Indeed, while for the *cis*-isomer the two dihedrals remain stable and close to the crystal structure value, in the *trans*-isomer case we observe a sudden reorientation of the aromatic ring of F334, which goes from 180° to 45°. This behavior is not unexpected as it can be related to the more rigid and more extended geometry of the *trans*-isomer that extends further from the buried binding pocket. This effect can also be pictorially appreciated in Figure 2E, in which the surface representing the solvent-excluded surface of F335 for the the *trans*-azobenzene-escitalopram can be seen extruding from the binding pocket and almost reaching the exterior of the protein. On the contrary, the *cis*-isomer behaves similar to the parent escitalopram. This may also be due to the rotation of F335, which is 45° for *trans*- and 180° for *cis*-azo-escitalopram The orientation of F335, as seen for *cis*-isomer, shields the orthosteric binding site, which, especially compared to the allosteric binding site, is buried deeper in the protein. F335 has recently been shown to play a crucial role in the occlusion of hSERT and thus in 5HT reuptake.^41^ Our preliminary observation point to a stronger perturbation of the outward-occluded protein conformation induced by the *trans*-isomer.

In order to assess the binding efficiency of the two isomers, we estimate the binding free energy difference between *cis*- and *trans*-azo-escitalopram. The free energy change between the unbound and bound *cis*/*trans*-ligand to the hSERT is calculated using the Molecular Mechanics / Poisson Boltzmann Surface Area (MM/PBSA) method, ^42,43^ a post-processing trajectory analysis technique, combined with the harmonic normal mode analysis method to approximate the entropic contribution. The enthalpic contribution (ΔH) is calculated for 2500 snapshots taken from the *cis*/*trans*-azo-escitalopram MD simulations between 1.1 *µ*s and 1.6 *µ*s, while the entropic (TΔS) contribution is estimated by using 50 snapshots (Section S3). The results, shown in Table 1, indicate a favorable binding for both isomers. As expected, the binding is largerly driven by a favorable enthalpic contribution, which overcomes an unfavorable entropic contribution, due to the reduction of the positional degrees of freedom upon binding. The binding of the two isomers provides a free energy gain of about 14 and 16 kcal mol^-1^for *cis*- and *trans*-isomers, respectively. This results suggests that both isomers may compete to occupy the binding pocket. The more favorable binding of the *trans*-isomer can be related to the smaller perturbation induced by the latter on the outward conformation of the protein. In particular, this is also coherent with the capacity of the *trans*-isomer to maintaining a more ideal binding pattern closer to the one exhibited by the parent escitalopram, which is instead more perturbed by the *cis*-isomer. On a side note, we see that the entropic contribution is almost the same for both isomers.

**Table 1:** Enthalpic (ΔH) and entropic (TΔS) contributions as well as resulting free energy (ΔG) calculated with the MM/PBSA method for *cis*/*trans*-azo-escitalopram. All numbers are given in kcal mol^-1^. ΔH and TΔS are given tog her with their standard error of mean *σ*_*H,S*_. The error of Δ*G* (*σ*_*G*_) is given by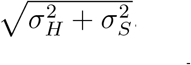

| | $\Delta H$ | $T\Delta S$ | $\Delta G$ |
| --- | --- | --- | --- |
| *cis* | -36.41 ± 0.11 | -22.38 ± 2.62 | -14.03 ± 2.62 |
| *trans* | -38.09 ± 0.09 | -22.10 ± 0.66 | -15.99 ± 0.67 |

While we have shown that both isomers may efficiently and favorably bind the hSERT, thus potentially inhibiting its activity, the reason behind the higher inhibiting potentiality of the *cis*-isomer remains elusive. Yet, it has been evidenced that inhibition of the parent escitalopram is due to the fact that its binding in the orthosteric pockets locks the hSERT in its outward-open conformation. Therefore, in the following, we analyze the structural descriptors that can be used to classify and differentiate between the hSERT conformations adopted during the uptake cycle (Fig. 1A).

As shown in Figure 3A, a channel connecting the cytoplasm with the extracellular medium may be identified. The latter also encompasses the orthosteric binding site and is delimited by two gates, the extracellular, formed by the I172, Y176, R104, E493, and F335 amino acids, and the intracellular gate defined by E444, R462, R79, D452, E80, Y350, and K275. The accessibility of the former gates may be related to the physiological relevant states of the transporter, namely the outward-open (cf. Fig. 1A) which allows the serotonin uptake and the outward-occluded, which instead leads to its internalization (cf. Fig. 1A). As pointed out by Gradisch et al.^44^ and pictorially represented in Figure 3B, the discrimination between these states can be achieved by monitoring the distance between the transmembrane TM6up *α−*helix and the TM6a and TM1b *α−*helices, respectively. Longer distances are characteristic of an outward-open conformation, while shorter distances point to an outward-occluded state. Interestingly, a significant difference is also observed upon binding of the natural 5HT substrate. 5HT binding leads to even shorter inter-helical distances, which favors the transitions towards inward-occluded and inward-open states, allowing the release of 5HT into the neuron, in line with the hSERT’s biological role.^41^ Figures 3C and D show the measured distribution of the extracellular gate and a time series of the inter-helical distances for hSERT bound to *trans*- and *cis*-azo-escitalopram. We clearly see that the *trans*-isomer is consistently giving shorter helical distances, both considering key residues (as part of the extracellular gate, see Fig. 3C) or the whole secondary structure (see Fig. 3D). As expected, the intracellular gate remains unchanged for both *cis*- and *trans*-azo-escitalopram (Fig. S1). Thus, it appears that the *trans*-isomer perturbs the structural descriptors pointing to shorter inter-helical distances. However, the *trans*-isomer shows a more flexible conformation of the transporter, with the TM6a-TM9up distance varying between the outward-open and outward-occluded references. In turn, the *cis*-isomer consistently shows distances closer to the outward-open conformation, which may be related to a more efficient locking of the hSERT in the purely outward-open state, thus inhibiting the transporter activity more efficiently.

**Figure 3.**
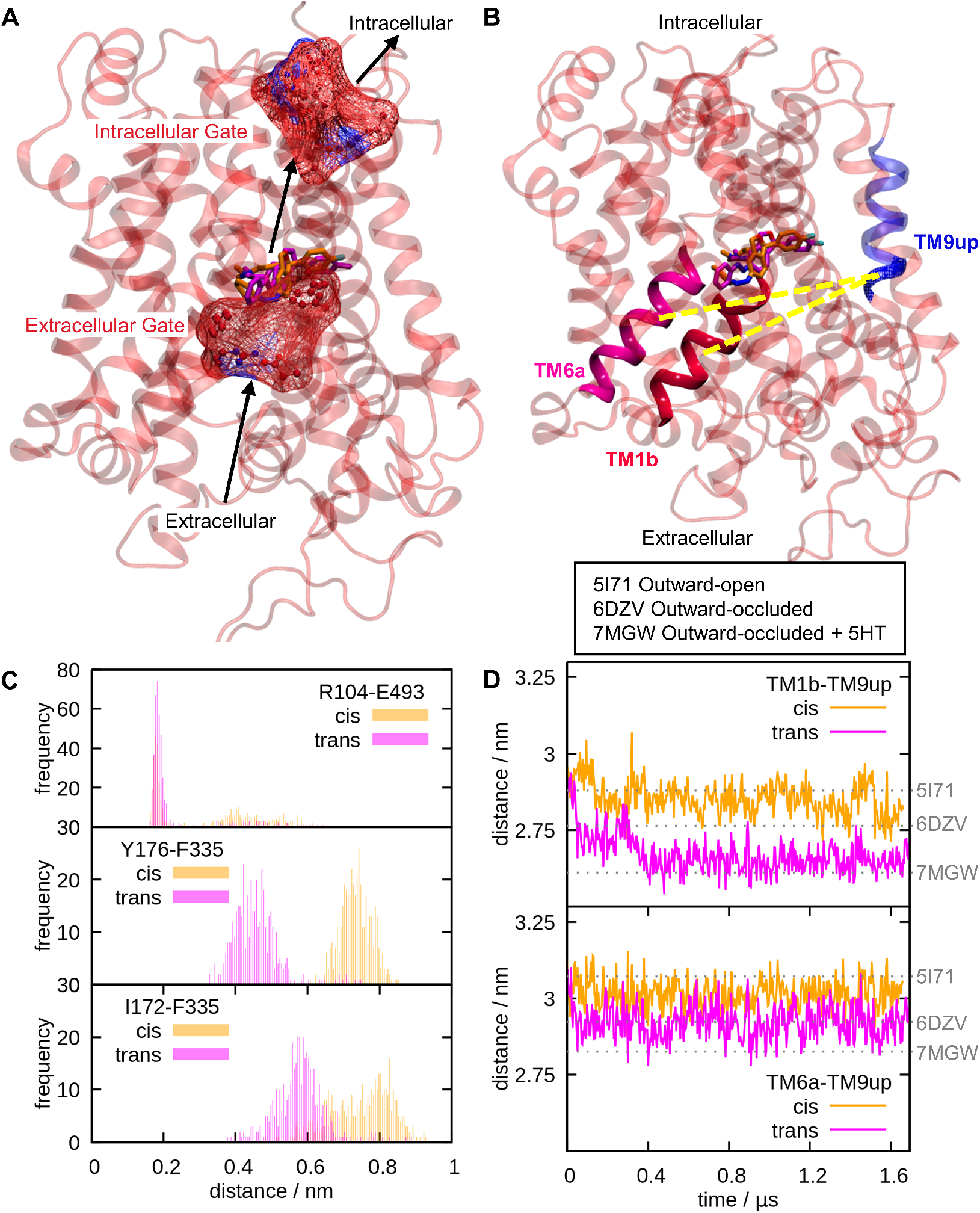
A) Location of the extra- and intracellular gates of hSERT. B) Transmembrane helices TM6a (pink, G324 to L337), TM1b (red, L99 to Q111), and TM9up (blue, F475 to S477) used to distinguish between the outward-open and outward-occluded hSERT conformation. PDB structures used as references are given. C) Histograms of three distances between amino acids (R104-E493, Y176-F335, I172-F335) involved in the extracellular gate of hSERT for *trans*- and *cis*-azo-escitalopram in pink and orange, respectively. D) Plots of the distances between TM1b and TM9up (top) and TM6a and TM9up (bottom) for *trans*- and *cis*-azo-escitalopram in pink and orange, respectively. The reference values (dotted lines) are given by the reference structures named by their PDB id (see also Fig. 3B).

In conclusion, by using long-scale MD simulation and free energy estimations, we have identified the molecular mechanisms leading to the inhibition of the transporting activity of the hSERT upon binding of azo-escitalopram, and its modulation induced by the action of light excitation. In particular, we have shown that while both isomers can favorably bind to the orthosteric pocket, their local arrangement is different and may strongly modulate the conformational transition cycle of the hSERT. Indeed, the *cis*-isomer appears to efficiently lock the hSERT in an outward-open conformation. Contrarily, the *trans*-isomer while providing shorter inter-helical distances also shows some instances of conformation transitions between outward-open and outward-occluded conformations. Thus, we hypothesize that *cis* is more efficient in maintaining the hSERT in the outward-open state compared to its *trans* counterpart. Even if this hypothesis is sounding, enhanced sampling MD simulations would be necessary to firmly establish it, in particular by exploring the free energy profiles for the outward-occluded to inward-occluded conformational transitions in the presence of both isomers. Such calculations would require a considerable computational effort, and an optimization of the chosen collective variable to reduce the dimensionality of the problem. Work along these lines can be the object of a forthcoming contribution.

Our results provide the first resolution at molecular level of the modulation of serotonin transporters by molecular photoswitches and evidence a delicate and subtle balance between structural and thermodynamic effects, which ultimately controls the global photoswitch inhibition capacity. As such, thris contribution may allow for a better and rational design of novel hSERT photo-activable inhibitors, which could also be profitable in optogenetics applications.

## Acknowledgement

N.K.S. and L.G. thank the Austrian Science Fund (FWF), W 1232 (MolTag) for financial support. A.M. thanks ANR and CGI for their financial support of this work through Labex SEAM ANR 11 LABX 086, ANR 11 IDEX 05 02. The support of the IdEx “Université Paris 2019” ANR-18-IDEX-0001 and of the Platform P3MB is gratefully acknowledged.

## Supporting Information Available

Computational details of DFT calculations and XYZ-coordinates of the optimized *trans*-/*cis-*azo-escitalopram. Computational details, results, and assessment of MD and MM/PBSA calculations.

