## Supplementary Information for "Molecular Photoswitches Regulating the Activity of the Human Serotonin Transporter"

### Contents

**S1 Quantum Chemical Calculations** **S4**

**S2 Molecular Dynamics Simulations** **S7**

**S3 MM/PBSA Calculations** **S9**

**References** **S10**

### List of Figures

S1 Histogram of the minimum distances between amino acids (E444, R462, R79, D452, E80, Y350, K275) involved in the intracellular gate of hSERT evolving over time of the MD simulations. . . . . . . . . . . . . . . . . . . . . . . . . . S7

S2 Time series of the distances of the center of masses (COM, top) and minimum distances (bottom) between amino acids involved in the extracellular (R104, E493, Y176, F335, Y172) and intracellular (E444, R462, R79, D452, E80, Y350, K275) gates of hSERT in the MD simulation of *cis*- (left) and *trans*-azo-escitalopram (right). . . . . . . . . . . . . . . . . . . . . . . . . . . . . . S8

### List of Tables

S1 Energies (in Hartree) of *cis*- and *trans*-azo-escitalopram optimized at the CAM-B3LYP-D3/def2-TZVP@SMD(DMSO) level of theory. . . . . . . . . . . S4

S2 XYZ coordinates of *cis*-azo-escitalopram optimized at the CAM-B3LYP-D3/def2-TZVP@SMD(DMSO) level of theory. . . . . . . . . . . . . . . . . . . . . . S5

S3 XYZ coordinates of *trans*-azo-escitalopram optimized at the CAM-B3LYP-D3/def2-TZVP@SMD(DMSO) level of theory. . . . . . . . . . . . . . . . . . . S6

S4 Enthalpic (ΔH) and entropic (TΔS) contributions calculated with MMPBSA for *cis*/*trans*-azo-escitalopram. All numbers are given in kcal $mol^{-1}$ together with their standard error of mean. . . . . . . . . . . . . . . . . . . . . . . . . S9

### S1 Quantum Chemical Calculations

In this section we provide the details of the quantum chemical optimization of the ground state geometry of *cis*- and *trans*-azo-escitalopram. After a preliminary conformational search done using CREST 2.10.2.,[1,2] we used density functional theory (DFT) with the CAM-B3LYP functional[3] with dispersion correction (D3)[4] and the def2-TZVP basis set[5,6] to optimize the ground state of both isomers. To simulating the experimental conditions, we opted to implicitly solvate the systems in dimethylsulfoxide (DMSO) by using the solvation model based on density (SMD) approach.[7] These calculations have been performed using Gaussian 16.[8] Gibbs free energies at 298.15 K and the optimized XYZ-coordinates are reported in Tables S1 to S3.

**Table S1: Energies (in Hartree) of *cis*- and *trans*-azo-escitalopram optimized at the CAM-B3LYP-D3/def2-TZVP@SMD(DMSO) level of theory.**

| Type of Energy | *cis* | *trans* |
|---|---|---|
| Sum of electronic and zero-point Energies | -1306.966839 | -1306.982939 |
| Sum of electronic and thermal Energies | -1306.940655 | -1306.956337 |
| Sum of electronic and thermal Enthalpies | -1306.939711 | -1306.955393 |
| Sum of electronic and thermal Free Energies | -1307.027302 | -1307.044852 |

**Table S2: XYZ coordinates of *cis*-azo-escitalopram optimized at the CAM-B3LYP-D3/def2-TZVP@SMD(DMSO) level of theory.**

| element | x | y | z |
|---|---|---|---|
| F | -0.69257 | 5.71322 | -1.09076 |
| C | -0.86121 | 4.44818 | -0.63861 |
| C | -1.30801 | 3.48593 | -1.51711 |
| C | -1.47715 | 2.19569 | -1.04728 |
| C | -1.20531 | 1.87087 | 0.27879 |
| C | -0.75818 | 2.87125 | 1.12999 |
| C | -0.58174 | 4.17103 | 0.67781 |
| H | -0.23395 | 4.95625 | 1.33548 |
| H | -0.54941 | 2.63775 | 2.16344 |
| C | -1.38710 | 0.43891 | 0.78208 |
| C | -0.37573 | -0.48361 | 0.13569 |
| C | -0.33262 | -1.00118 | -1.14502 |
| C | 0.71695 | -1.83672 | -1.49015 |
| C | 1.72591 | -2.10296 | -0.57370 |
| N | 2.75164 | -3.01563 | -0.97956 |
| N | 3.95096 | -2.75835 | -0.86608 |
| C | 4.43117 | -1.47326 | -0.45511 |
| C | 4.10253 | -0.32001 | -1.15437 |
| C | 4.66934 | 0.88506 | -0.77850 |
| C | 5.54711 | 0.94236 | 0.29373 |
| C | 5.88070 | -0.21793 | 0.97614 |
| C | 5.34176 | -1.43255 | 0.58890 |
| H | 5.60831 | -2.34842 | 1.10073 |
| H | 6.57293 | -0.17978 | 1.80719 |
| H | 5.98013 | 1.88894 | 0.58945 |
| H | 4.41947 | 1.78531 | -1.32482 |
| H | 3.41394 | -0.36916 | -1.98682 |
| C | 1.67056 | -1.60590 | 0.72266 |
| C | 0.60015 | -0.80539 | 1.05904 |
| C | 0.26822 | -0.14690 | 2.35818 |
| H | 0.96722 | 0.66618 | 2.58493 |
| H | 0.26249 | -0.83946 | 3.20123 |
| O | -1.05192 | 0.35884 | 2.17373 |
| H | 2.43973 | -1.85211 | 1.44194 |
| H | 0.76933 | -2.27838 | -2.47674 |
| H | -1.10942 | -0.78104 | -1.86469 |
| C | -2.83419 | -0.01454 | 0.60923 |
| C | -3.10750 | -1.42034 | 1.11509 |
| C | -4.55692 | -1.84152 | 0.93237 |
| N | -5.03569 | -1.75606 | -0.44027 |
| C | -6.45795 | -2.02129 | -0.50282 |
| H | -6.71077 | -3.04405 | -0.17527 |
| H | -6.81711 | -1.89804 | -1.52547 |
| H | -6.99788 | -1.32073 | 0.13578 |
| C | -4.30881 | -2.63147 | -1.33712 |
| H | -3.25699 | -2.35187 | -1.38695 |
| H | -4.72288 | -2.54993 | -2.34267 |
| H | -4.36748 | -3.68918 | -1.02882 |
| H | -4.68223 | -2.86775 | 1.32039 |
| H | -5.19384 | -1.19402 | 1.54021 |
| H | -2.44526 | -2.13113 | 0.61813 |
| H | -2.87312 | -1.47682 | 2.17896 |
| H | -3.47550 | 0.69995 | 1.13229 |
| H | -3.08755 | 0.05325 | -0.44863 |
| H | -1.82485 | 1.43667 | -1.73437 |
| H | -1.51700 | 3.74603 | -2.54596 |

**Table S3: XYZ coordinates of *trans*-azo-escitalopram optimized at the CAM-B3LYP-D3/def2-TZVP@SMD(DMSO) level of theory.**

| element | x | y | z |
|---|---|---|---|
| N | -6.03562 | -2.39386 | -0.61919 |
| N | 4.65069 | -0.31880 | -0.55712 |
| C | -1.66522 | -0.11324 | 0.45061 |
| C | -2.42055 | -1.20154 | -0.30882 |
| C | -0.17110 | -0.17995 | 0.22064 |
| C | 0.47538 | -0.21139 | 1.43965 |
| C | -3.91140 | -1.23582 | -0.01403 |
| C | -2.19906 | 1.28327 | 0.13314 |
| C | -0.54638 | -0.18417 | 2.52814 |
| C | -4.58293 | -2.40328 | -0.71603 |
| C | 0.54769 | -0.21134 | -0.96405 |
| C | 1.85179 | -0.25943 | 1.51058 |
| C | -2.24236 | 1.74069 | -1.18066 |
| C | -2.64684 | 2.12050 | 1.14432 |
| C | 1.92712 | -0.26419 | -0.90897 |
| C | 2.57839 | -0.28296 | 0.32575 |
| C | -2.71799 | 3.00398 | -1.48286 |
| C | -3.12671 | 3.39134 | 0.86190 |
| C | -3.15045 | 3.80424 | -0.44835 |
| C | -6.50139 | -2.58519 | 0.74091 |
| C | -6.61178 | -3.39428 | -1.49599 |
| O | -1.78676 | -0.38603 | 1.85351 |
| C | 6.05781 | -0.37669 | -0.39362 |
| N | 3.98515 | -0.33824 | 0.48381 |
| C | 6.70521 | -0.45088 | 0.83801 |
| C | 8.08409 | -0.50353 | 0.87768 |
| C | 8.82384 | -0.48297 | -0.29940 |
| C | 8.17853 | -0.40915 | -1.52236 |
| C | 6.79548 | -0.35575 | -1.56942 |
| F | -3.61851 | 5.04158 | -0.73571 |
| H | -2.25268 | -1.05905 | -1.37832 |
| H | -1.96454 | -2.15688 | -0.03937 |
| H | -4.05571 | -1.30135 | 1.06443 |
| H | -4.38145 | -0.30756 | -0.34696 |
| H | -0.41514 | -0.98104 | 3.26136 |
| H | -0.54373 | 0.77516 | 3.05796 |
| H | -4.32598 | -2.36782 | -1.77757 |
| H | -4.17415 | -3.35150 | -0.32304 |
| H | 0.04387 | -0.20079 | -1.92121 |
| H | 2.37543 | -0.28490 | 2.45774 |
| H | -1.90435 | 1.10758 | -1.98941 |
| H | -2.63245 | 1.77649 | 2.16779 |
| H | 2.50941 | -0.29588 | -1.81818 |
| H | -2.75695 | 3.36600 | -2.50130 |
| H | -3.47927 | 4.04937 | 1.64465 |
| H | -6.16202 | -1.77734 | 1.38712 |
| H | -6.15054 | -3.53882 | 1.17081 |
| H | -7.59182 | -2.58933 | 0.75723 |
| H | -6.30825 | -4.41880 | -1.22225 |
| H | -6.30090 | -3.21424 | -2.52590 |
| H | -7.70068 | -3.34471 | -1.45399 |
| H | 6.12589 | -0.46678 | 1.74981 |
| H | 8.59125 | -0.56135 | 1.83213 |
| H | 9.90461 | -0.52484 | -0.25823 |
| H | 8.75099 | -0.39304 | -2.44044 |
| H | 6.27141 | -0.29769 | -2.51467 |

#### S2 Molecular Dynamics Simulations

Here we provide supporting figures that illustrate the MD simulations. A distribution plot of the distances involved in the extracellular gate was provided in the main manuscript in Figure 3C; a similar plot for the intracellular gate is provided in Figure S1. Further, Figure S2 shows the the time series of all the extra- and intracellular gate distances.

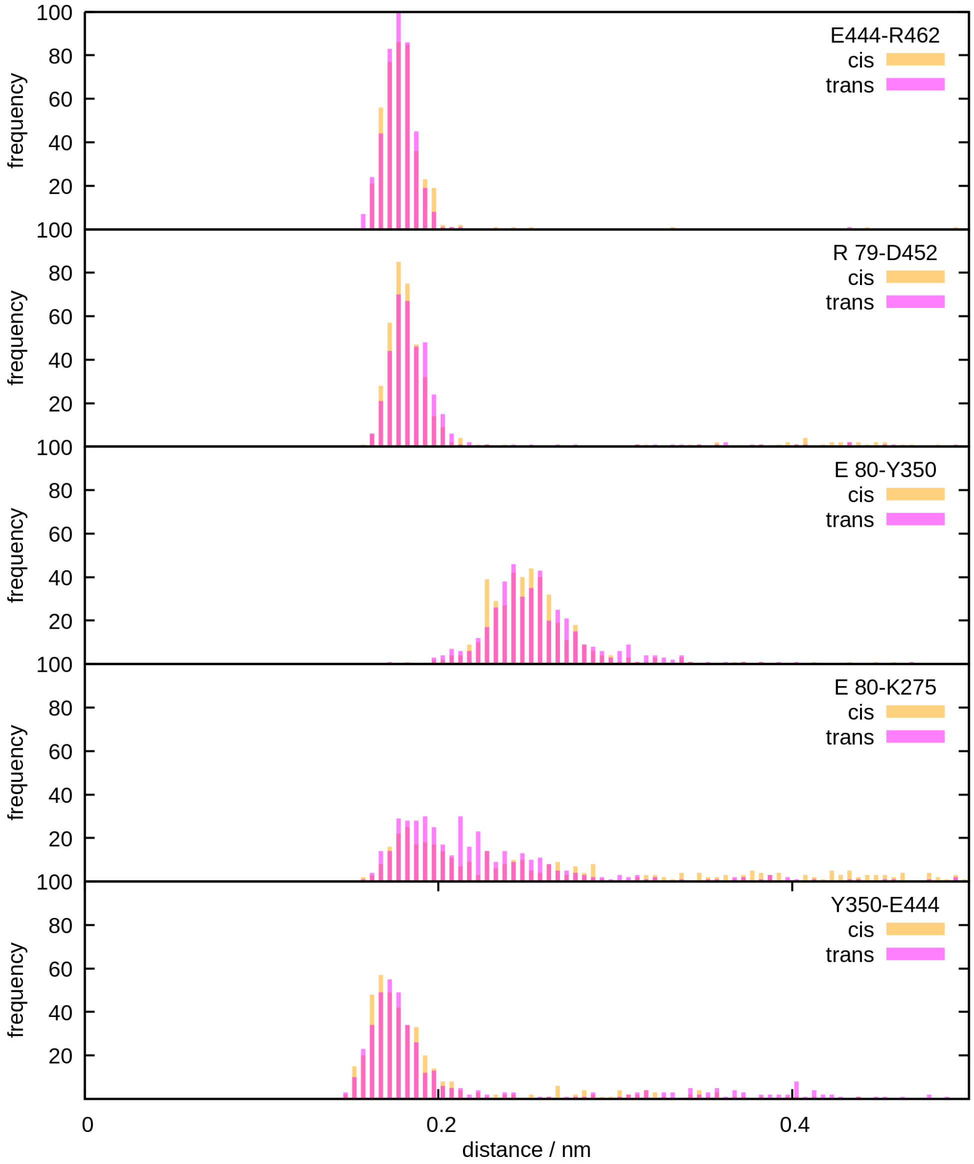


Figure S1: Histogram of the minimum distances between amino acids (E444, R462, R79, D452, E80, Y350, K275) involved in the intracellular gate of hSERT evolving over time of the MD simulations.

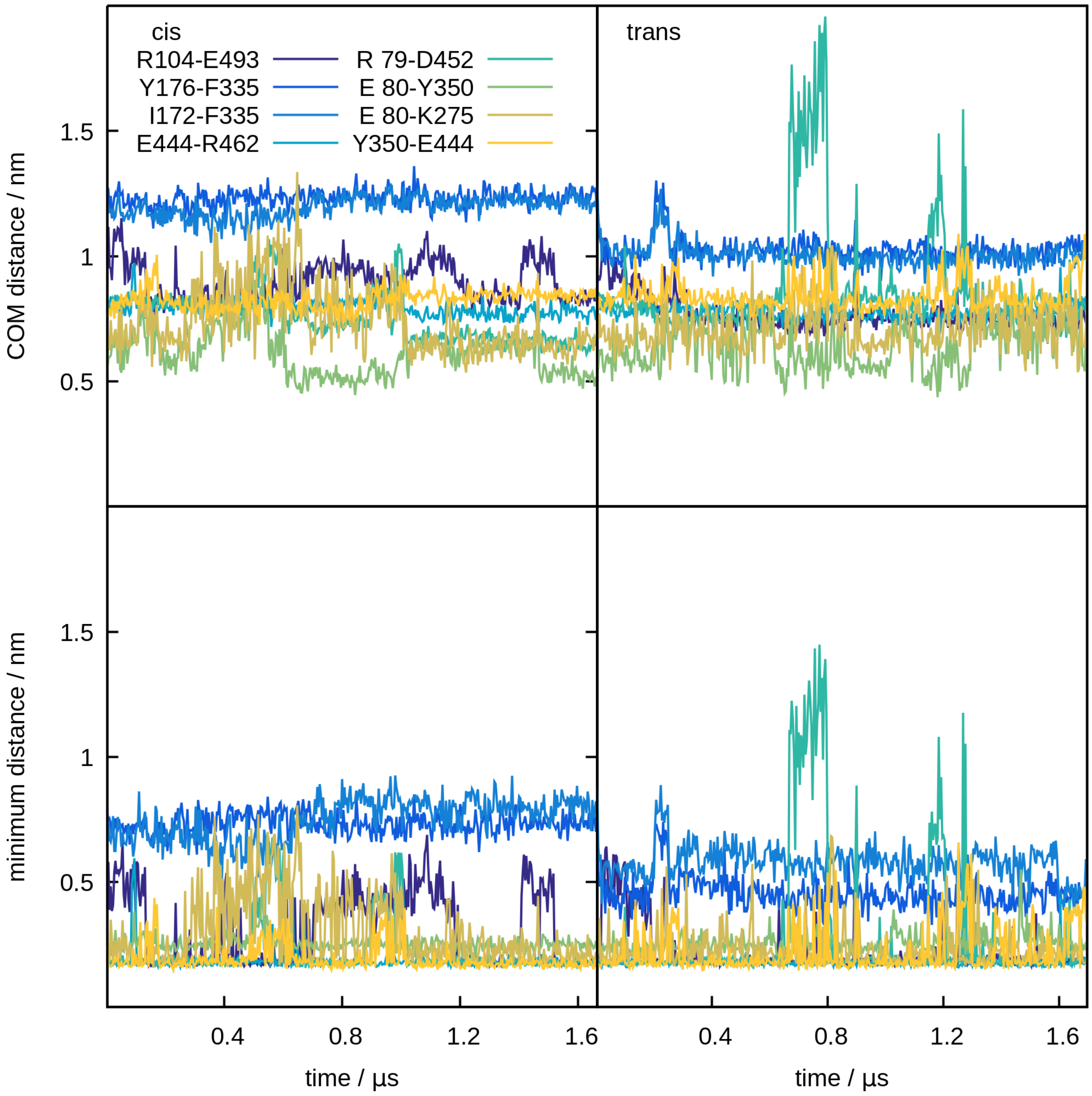


Figure S2: Time series of the distances of the center of masses (COM, top) and minimum distances (bottom) between amino acids involved in the extracellular (R104, E493, Y176, F335, Y172) and intracellular (E444, R462, R79, D452, E80, Y350, K275) gates of hSERT in the MD simulation of *cis*- (left) and *trans*-azo-escitalopram (right).

#### S3 MM/PBSA Calculations

The free energy change between the unbound and bound *cis*/*trans*-ligand to the hSERT is calculated using the Molecular Mechanics / Poisson Boltzmann Surface Area (MM/PBSA) method,[9,10] a post-processing trajectory analysis technique. The method is computationally inexpensive but assumes that the configurational space explored by the receptor and ligand is unchanged between the bound and unbound states. The MM/PBSA analysis was performed using the MMPBSA.py script[11] implemented in AMBER20[12] for 500 and 2500 snapshots taken from the *cis*/*trans*-azo-escitalopram MD simulations between 1.1 $\mu$s and 1.6 $\mu$s. The MMPBSA.py script was run with input values inspired by the "sample input for MMPBSA with membrane proteins" given in the ABER20 manual.[12] The membrane thickness was set to 36 Å, it was centered to the center of the protein, and a dielectric constant of 7 was assigned to the membrane. The Periodic Incomplete Cholesky Conjugate Gradient (PICCG) iterative solver was chosen and the total electrostatic energy and forces were computed with the particle-particle particle-mesh (P3M) procedure.[13] The atom-based cutoff distance for van der Waals interactions was set to 99 Åand the atom-based cutoff distance to remove short-range finite-difference interactions as well as to add pairwise charge-based interactions was set to 7 Å. The resulting enthalpy ($\Delta$H) is given in Table S4. The entropic contribution (T$\Delta$S = Temperature T $\times$ Entropy $\Delta$S) is estimated using an harmonic approximation, the normal mode analysis (NMA), with the same program. The results for 5, 20 and 50 snapshots from the *cis*/*trans*-azo-escitalopram MD simulations between 1.1 $\mu$s and 1.6 $\mu$s are given in Table S4.

**Table S4: Enthalpic ($\Delta$H) and entropic (T$\Delta$S) contributions calculated with MMPBSA for *cis*/*trans*-azo-escitalopram. All numbers are given in kcal mol$^{-1}$ together with their standard error of mean.**

| | $\Delta$H | | T$\Delta$S | | |
|---|---|---|---|---|---|
| # of snapshots | 500 | 2500 | 5 | 20 | 50 |
| *cis* | -36.39 ± 0.25 | -36.41 ± 0.11 | -18.90 ± 5.32 | -21.97 ± 1.01 | -22.38 ± 2.62 |
| *trans* | -38.02 ± 0.19 | -38.09 ± 0.09 | -17.23 ± 1.25 | -19.91 ± 0.87 | -22.10 ± 0.66 |

(8) Frisch, M. J.; Trucks, G. W.; Schlegel, H. B.; Scuseria, G. E.; Robb, M. A.; Cheese-

man, J. R.; Scalmani, G.; Barone, V.; Petersson, G. A.; Nakatsuji, H. et al. Gaussian16 Revision C.01. Gaussian Inc. Wallingford CT, 2016.
